# QSAR Model Based Gradient Boosting Regression of N-Arylsulfonyl-Indole-2-Carboxamide Derivatives as Inhibitors for Fructose-1,6-Bisphosphatase

**DOI:** 10.1101/2021.08.10.455890

**Authors:** Ziyi Zhao, Jialong Yang, Bowen Li, Tingting Sun, Hongzong Si, Tongshang Ni

## Abstract

As is known to all, diabetes metellius is a global health threaten and it has caused worldwide attention of scientists. To get a better investigation of the drug design of diabetes, we used heuristic method to established the linear model and used Gradient Boosting Regression to establish the nonlinear model of Fructose-1,6-Bisphosphatse inhibitor successively. In this study, 84 derivatives of N-Arylsulfonyl-Indole-2-Carboxamide were introduced into the models, two outstanding QSAR models with 2 molecule descriptors were established successfully. Grandient Boosting Regression rendered a good correlation with R^2^ of 0.943 and MSE of 0.135 for the training set, 0.916 and 0.213 for test set, which also proves the feasibility of the implementation of the new method GBR in the field of QSAR. Meanwhile, the optimal model displayed wonderful statistical significance. This study shows unlimited potential for design of new drugs for diabetes.

## 1. Introduction

Characterized by hyperglycemia, diabetes metellius (DM) is a chronic metabolic disease. It can do severe harm to the kidney, blood vessels, eyes, nerves and hearts. In addition, DM also threatens the safety of babies and pregnant women since it is associated with preterm delivery, birthweight extremes and congenital anomaly [1]. According to the statistics coming from the <u>WHO webpage</u>, 6 percentages of the population was diagnosed with DM. DM can be divided into type 1 diabetes metellius (T1DM) and type 2 diabetes metellius (T2DM), T2DM accounts for 90% of diagnosed DM. The main feature of T2DM is insulin resistance, leading to the higher risks of ischemic heart diseases and stroke [2]. At present, Hypoglycemic drugs mainly include biguanide drugs and sulfonylurea hypoglycemic drugs, the pharmacological mechanism of the majority of anti-diabetic drugs is to increase the secretion of insulin or avoid the insulin resistance[3]. However, sulfonylurea hypoglycemic drugs may cause hypoglycemia, weight gain and the typical side effect of biguanide drugs is gastrointestinal reactions[4]. Due to the complication caused by the conventional drugs, global attention has been focused on the development of novel drugs. As a consequence, a potential theory to put T2DM under control is of great medical significance.

Gluconeogensis (GNG) is the main endogenous glucose production process for providing glucose in liver and kidney[5], which takes on an important position in the onset of T2DM. Serving as an important rate-limiting enzyme in the GNG pathway, fructose-1,6-bisphosphatse(FBPase) catalyze irreversible reaction from fructose-1,6-bisphosphate to fructose-6-phosphate[6]. FBPase has two isoforms which exist in liver and muscle, respectively. According to the previous researches, not only can it participate in the energy metabolism and glucose homeostasis, but also it interacts with mitochondrial and nuclear proteins[7]. Thus, there is no denying that FBPase is a promising and attractive target to affect the GNG pathway and control the level of blood glucose, N-Arylsulfonyl-Indole-2-Carboxamide Derivatives shows great research values Inhibitors for Fructose-1,6-Bisphosphatase.

Computer-aided drug discovery (CADD) is a method of designing and optimizing pilot compounds through computer calculation, stimulation and budgeting of the relationship between biomolecules and drugs[8]. Quantitative-structure-activity relationship (QSAR) is one of the most widely used methods in CADD. This approach established the quantitative relationship between physiological activity or certain properties of a series of compounds and their physical or chemical properties through some mathematical statistical models. We can predict the activity of new compounds with high predicted ability with these models[9]. To study the inhibitory effect of N-Arylsulfonyl-Indole-2-Carboxamide Derivatives, we build up two models with gradient boosting regression (GBR) and heuristic method(HM). As far as we know, these derivatives has been designed and synthesised as potent, selective, and orally bioavailable FBPase inhibitors in the recent study, at the meantime, several promising candidates have been chosen as human liver FBPase for its high inhibitory activity through structure-activity relationship studies. Additional in-depth studies of these novel compounds will be needed to fully characterize their roles as FBPase inhibitors and we are not aware of any publications with N-Arylsulfonyl-Indole-2-Carboxamide derivatives on the QSAR model based on GBR. This work promises a wonderful prospect to the further studies of T2DM.

## 2. Method

### 2.1 Data

The inhibitory data of the enzyme FBPase of 84 compounds were selected from the literature[10]. The screening criteria were dismissing the derivatives without the accurate inhibitory data. The biological activities were expressed by the half-maximum inhibitory concentration (IC_50_) values, the structure, experimental and predictive IC_50_ values of the derivatives were listed in the Tab.1. Then, we normalized the data to decrease the impact of the dimension for getting a global optimum. Thus, we used the square root of the IC_50_ to process the data, 84 compounds were randomly divided into 21 compounds of test set and 63 compounds of training set, respectively. Training set is the data sample used for model fitting while test set is the sample set aside separately during the model training process, which can be used to make a preliminary assessment of the model’s predicative ability. In the Tab.1,the text set was marked with *.

**Table 1.**
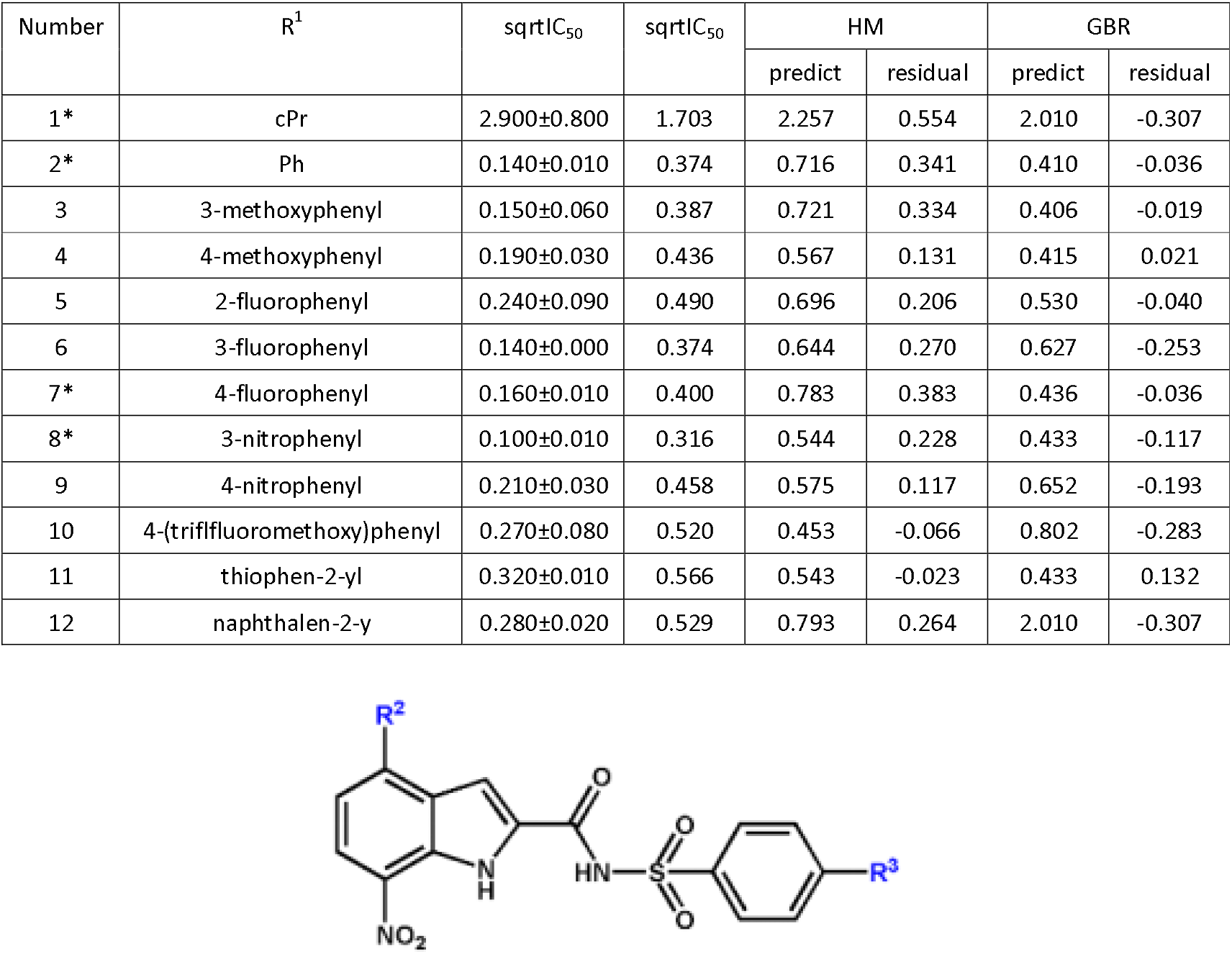
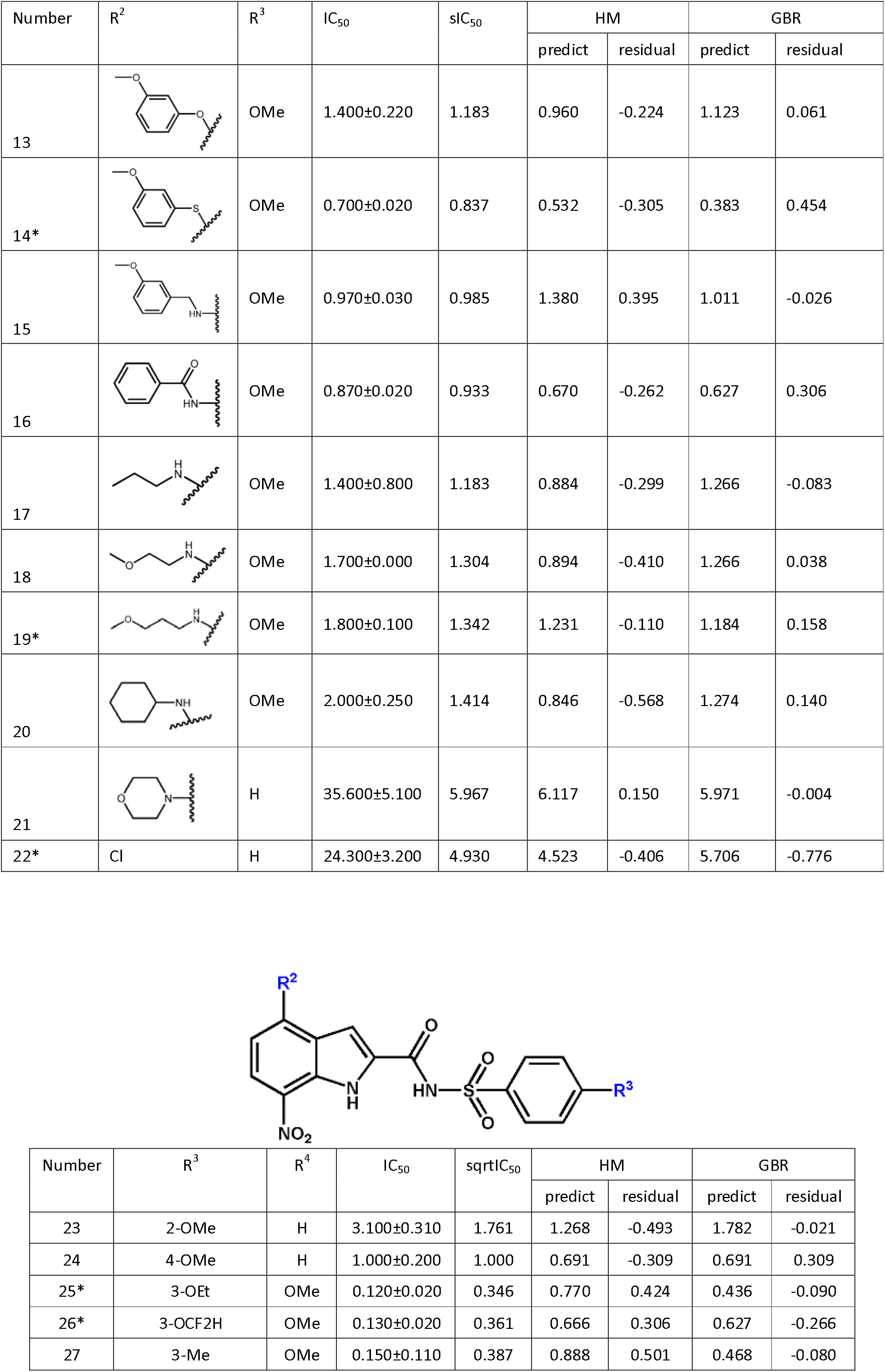
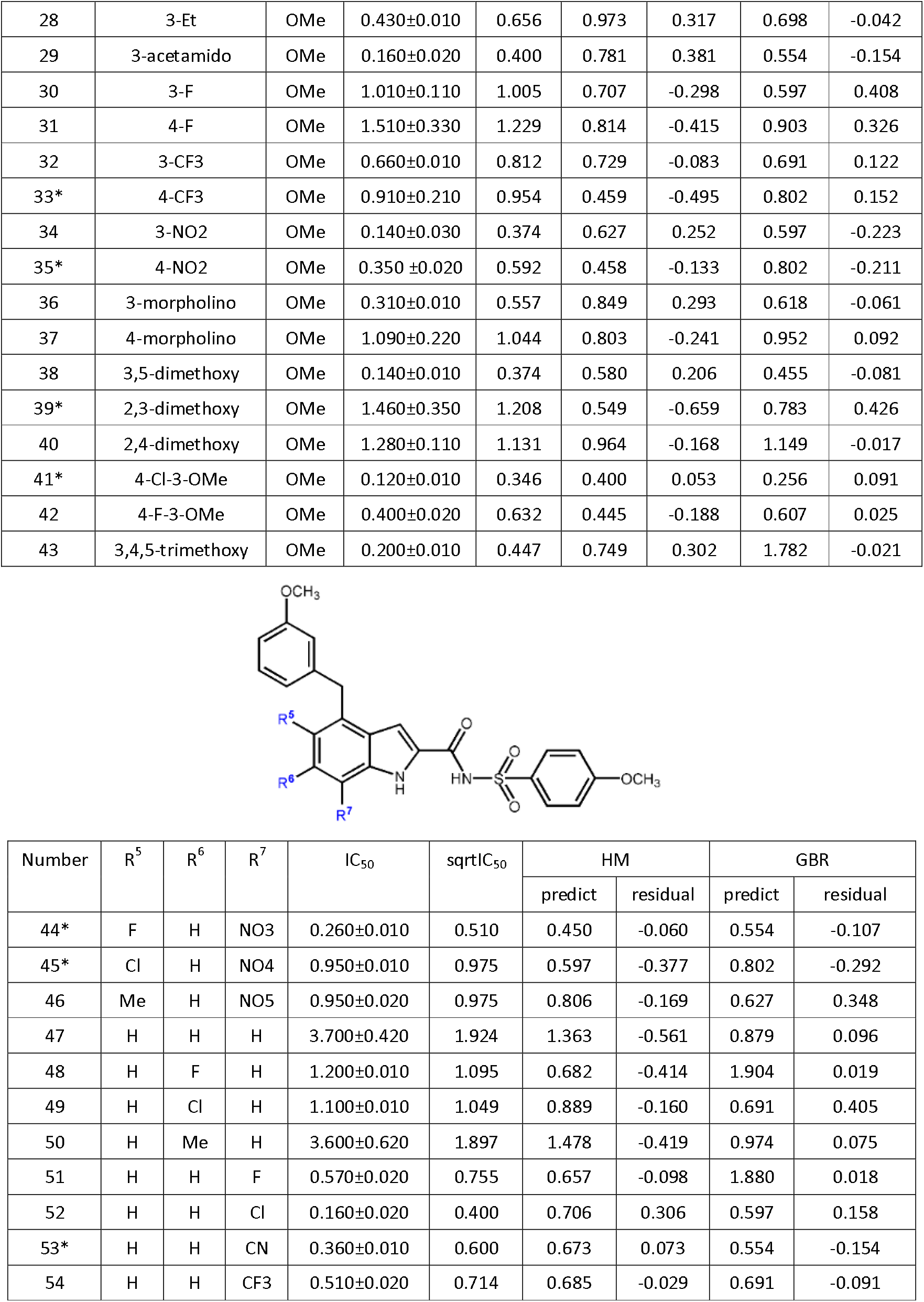
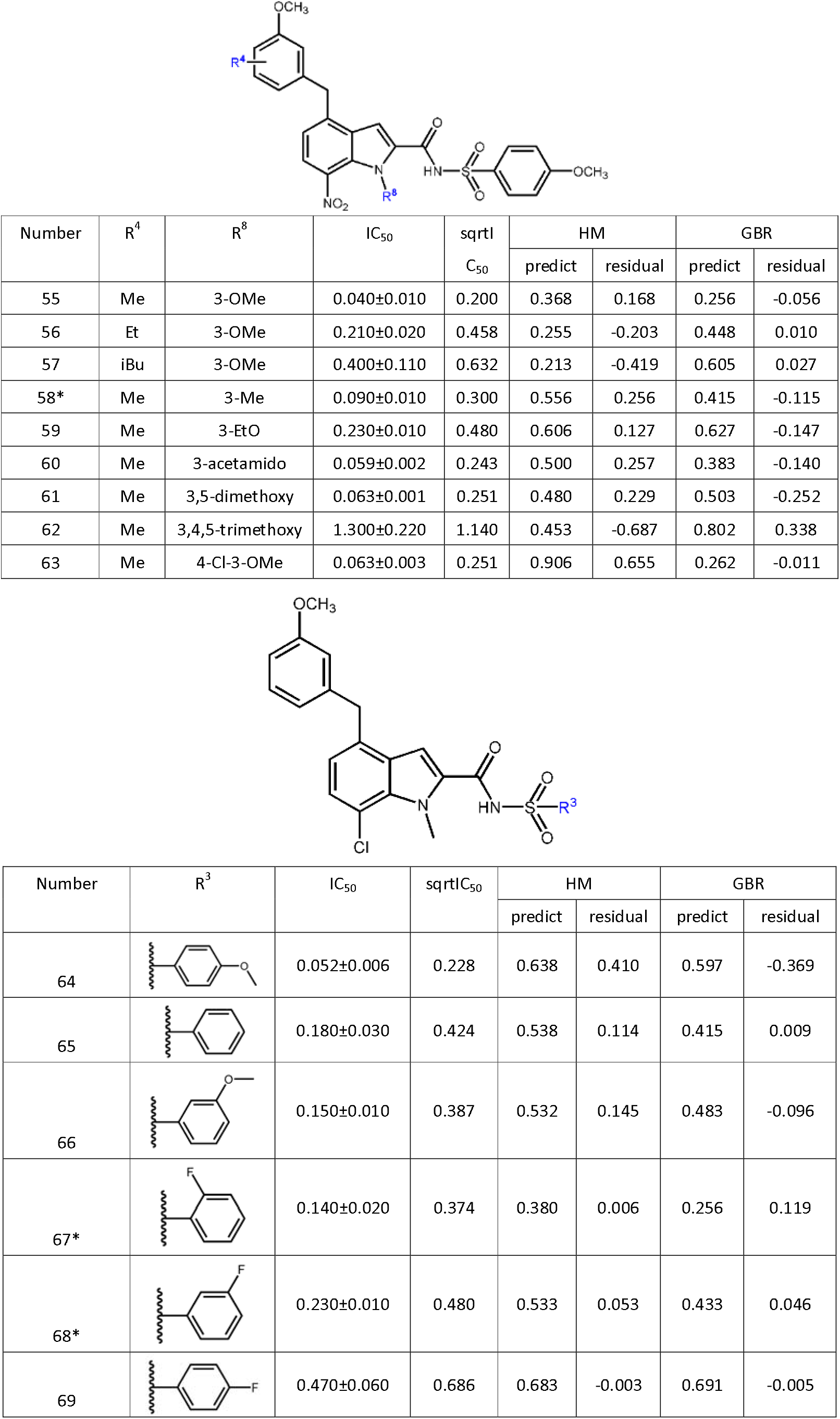
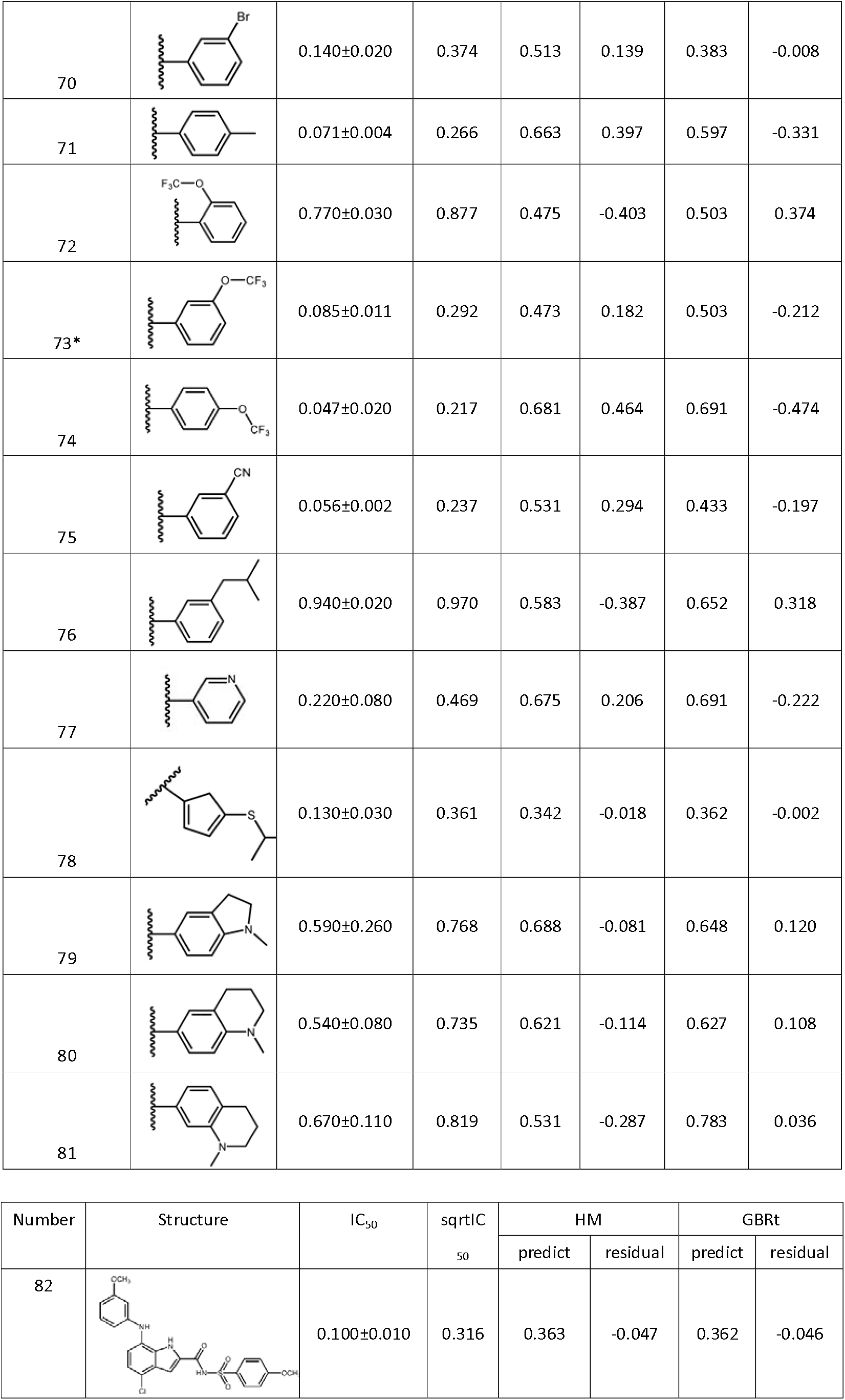
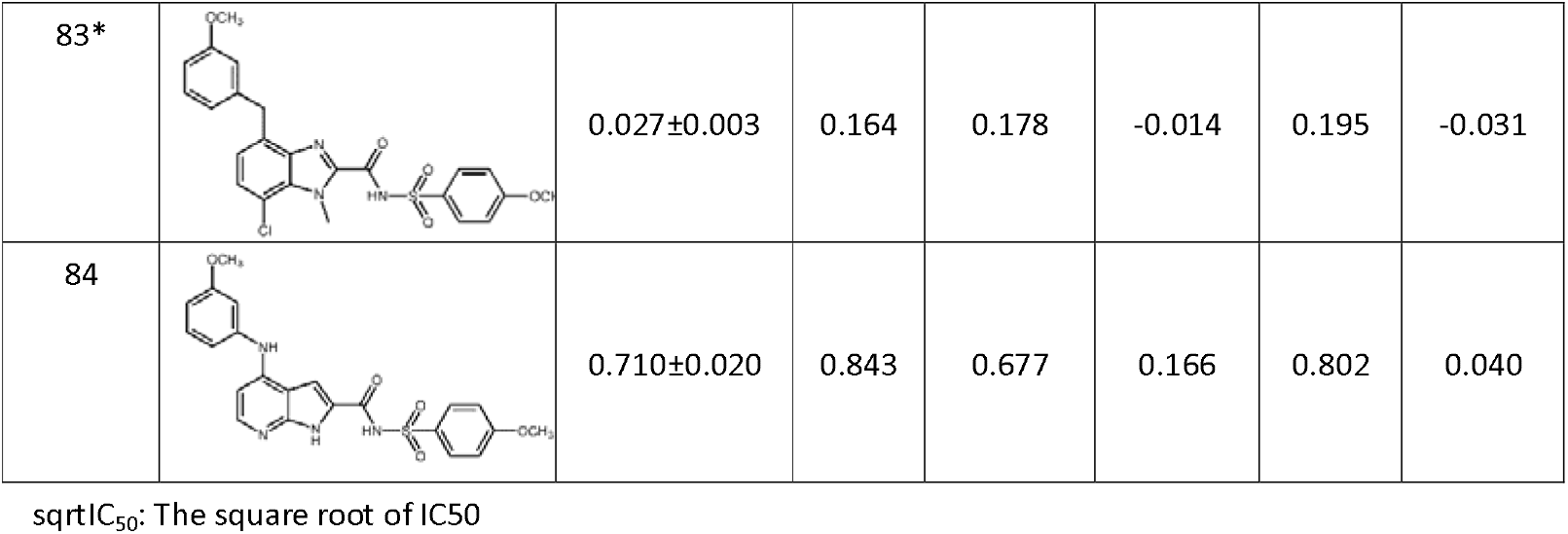

| Number | R<sup>1</sup> | sqrt(C<sub>50</sub>) | sqrt(C<sub>50</sub>) | HM |  | GBR |  |
| --- | --- | --- | --- | --- | --- | --- | --- |
|  |  |  |  | predict | residual | predict | residual |
| 1* | cPr | 2.900±0.800 | 1.703 | 2.257 | 0.554 | 2.010 | -0.307 |
| 2* | Ph | 0.140±0.010 | 0.374 | 0.716 | 0.341 | 0.410 | -0.036 |
| 3 | 3-methoxyphenyl | 0.150±0.060 | 0.387 | 0.721 | 0.334 | 0.406 | -0.019 |
| 4 | 4-methoxyphenyl | 0.190±0.030 | 0.436 | 0.567 | 0.131 | 0.415 | 0.021 |
| 5 | 2-fluorophenyl | 0.240±0.090 | 0.490 | 0.696 | 0.206 | 0.530 | -0.040 |
| 6 | 3-fluorophenyl | 0.140±0.000 | 0.374 | 0.644 | 0.270 | 0.627 | -0.253 |
| 7* | 4-fluorophenyl | 0.160±0.010 | 0.400 | 0.783 | 0.383 | 0.436 | -0.036 |
| 8* | 3-nitrophenyl | 0.100±0.010 | 0.316 | 0.544 | 0.228 | 0.433 | -0.117 |
| 9 | 4-nitrophenyl | 0.210±0.030 | 0.458 | 0.575 | 0.117 | 0.652 | -0.193 |
| 10 | 4-(trifluoromethoxy)phenyl | 0.270±0.080 | 0.520 | 0.453 | -0.066 | 0.802 | -0.283 |
| 11 | thiophen-2-yl | 0.320±0.010 | 0.566 | 0.543 | -0.023 | 0.433 | 0.132 |
| 12 | naphthalen-2-yl | 0.280±0.020 | 0.529 | 0.793 | 0.264 | 2.010 | -0.307 |

| Number | R<sup>2</sup> | R<sup>3</sup> | IC<sub>50</sub> | sIC<sub>50</sub> | HM |  | GBR |  |
| --- | --- | --- | --- | --- | --- | --- | --- | --- |
|  |  |  |  |  | predict | residual | predict | residual |
| 13 |  | OMe | 1.400±0.220 | 1.183 | 0.960 | -0.224 | 1.123 | 0.061 |
| 14* |  | OMe | 0.700±0.020 | 0.837 | 0.532 | -0.305 | 0.383 | 0.454 |
| 15 |  | OMe | 0.970±0.030 | 0.985 | 1.380 | 0.395 | 1.011 | -0.026 |
| 16 |  | OMe | 0.870±0.020 | 0.933 | 0.670 | -0.262 | 0.627 | 0.306 |
| 17 |  | OMe | 1.400±0.800 | 1.183 | 0.884 | -0.299 | 1.266 | -0.083 |
| 18 |  | OMe | 1.700±0.000 | 1.304 | 0.894 | -0.410 | 1.266 | 0.038 |
| 19* |  | OMe | 1.800±0.100 | 1.342 | 1.231 | -0.110 | 1.184 | 0.158 |
| 20 |  | OMe | 2.000±0.250 | 1.414 | 0.846 | -0.568 | 1.274 | 0.140 |
| 21 |  | H | 35.600±5.100 | 5.967 | 6.117 | 0.150 | 5.971 | -0.004 |
| 22* | Cl | H | 24.300±3.200 | 4.930 | 4.523 | -0.406 | 5.706 | -0.776 |

| Number | R<sup>3</sup> | R<sup>4</sup> | IC<sub>50</sub> | sqrt(IC<sub>50</sub>) | HM |  | GBR |  |
| --- | --- | --- | --- | --- | --- | --- | --- | --- |
|  |  |  |  |  | predict | residual | predict | residual |
| 23 | 2-OMe | H | 3.100±0.310 | 1.761 | 1.268 | -0.493 | 1.782 | -0.021 |
| 24 | 4-OMe | H | 1.000±0.200 | 1.000 | 0.691 | -0.309 | 0.691 | 0.309 |
| 25* | 3-OEt | OMe | 0.120±0.020 | 0.346 | 0.770 | 0.424 | 0.436 | -0.090 |
| 26* | 3-OCF<sub>2</sub>H | OMe | 0.130±0.020 | 0.361 | 0.666 | 0.306 | 0.627 | -0.266 |
| 27 | 3-Me | OMe | 0.150±0.110 | 0.387 | 0.888 | 0.501 | 0.468 | -0.080 |
| 28 | 3-Et | OMe | 0.430±0.010 | 0.656 | 0.973 | 0.317 | 0.698 | -0.042 |
| 29 | 3-acetamido | OMe | 0.160±0.020 | 0.400 | 0.781 | 0.381 | 0.554 | -0.154 |
| 30 | 3-F | OMe | 1.010±0.110 | 1.005 | 0.707 | -0.298 | 0.597 | 0.408 |
| 31 | 4-F | OMe | 1.510±0.330 | 1.229 | 0.814 | -0.415 | 0.903 | 0.326 |
| 32 | 3-CF<sub>3</sub> | OMe | 0.660±0.010 | 0.812 | 0.729 | -0.083 | 0.691 | 0.122 |
| 33* | 4-CF<sub>3</sub> | OMe | 0.910±0.210 | 0.954 | 0.459 | -0.495 | 0.802 | 0.152 |
| 34 | 3-NO<sub>2</sub> | OMe | 0.140±0.030 | 0.374 | 0.627 | 0.252 | 0.597 | -0.223 |
| 35* | 4-NO<sub>2</sub> | OMe | 0.350 ±0.020 | 0.592 | 0.458 | -0.133 | 0.802 | -0.211 |
| 36 | 3-morpholino | OMe | 0.310±0.010 | 0.557 | 0.849 | 0.293 | 0.618 | -0.061 |
| 37 | 4-morpholino | OMe | 1.090±0.220 | 1.044 | 0.803 | -0.241 | 0.952 | 0.092 |
| 38 | 3,5-dimethoxy | OMe | 0.140±0.010 | 0.374 | 0.580 | 0.206 | 0.455 | -0.081 |
| 39* | 2,3-dimethoxy | OMe | 1.460±0.350 | 1.208 | 0.549 | -0.659 | 0.783 | 0.426 |
| 40 | 2,4-dimethoxy | OMe | 1.280±0.110 | 1.131 | 0.964 | -0.168 | 1.149 | -0.017 |
| 41* | 4-Cl-3-OMe | OMe | 0.120±0.010 | 0.346 | 0.400 | 0.053 | 0.256 | 0.091 |
| 42 | 4-F-3-OMe | OMe | 0.400±0.020 | 0.632 | 0.445 | -0.188 | 0.607 | 0.025 |
| 43 | 3,4,5-trimethoxy | OMe | 0.200±0.010 | 0.447 | 0.749 | 0.302 | 1.782 | -0.021 |

| Number | R<sup>5</sup> | R<sup>6</sup> | R<sup>7</sup> | IC<sub>50</sub> | sqrtIC<sub>50</sub> | HM |  | GBR |  |
| --- | --- | --- | --- | --- | --- | --- | --- | --- | --- |
|  |  |  |  |  |  | predict | residual | predict | residual |
| 44* | F | H | NO3 | 0.260±0.010 | 0.510 | 0.450 | -0.060 | 0.554 | -0.107 |
| 45* | Cl | H | NO4 | 0.950±0.010 | 0.975 | 0.597 | -0.377 | 0.802 | -0.292 |
| 46 | Me | H | NO5 | 0.950±0.020 | 0.975 | 0.806 | -0.169 | 0.627 | 0.348 |
| 47 | H | H | H | 3.700±0.420 | 1.924 | 1.363 | -0.561 | 0.879 | 0.096 |
| 48 | H | F | H | 1.200±0.010 | 1.095 | 0.682 | -0.414 | 1.904 | 0.019 |
| 49 | H | Cl | H | 1.100±0.010 | 1.049 | 0.889 | -0.160 | 0.691 | 0.405 |
| 50 | H | Me | H | 3.600±0.620 | 1.897 | 1.478 | -0.419 | 0.974 | 0.075 |
| 51 | H | H | F | 0.570±0.020 | 0.755 | 0.657 | -0.098 | 1.880 | 0.018 |
| 52 | H | H | Cl | 0.160±0.020 | 0.400 | 0.706 | 0.306 | 0.597 | 0.158 |
| 53* | H | H | CN | 0.360±0.010 | 0.600 | 0.673 | 0.073 | 0.554 | -0.154 |
| 54 | H | H | CF<sub>3</sub> | 0.510±0.020 | 0.714 | 0.685 | -0.029 | 0.691 | -0.091 |

| Number | R<sup>4</sup> | R<sup>8</sup> | IC<sub>50</sub> | sqrtIC<sub>50</sub> | HM |  | GBR |  |
| --- | --- | --- | --- | --- | --- | --- | --- | --- |
|  |  |  |  |  | predict | residual | predict | residual |
| 55 | Me | 3-OMe | 0.040±0.010 | 0.200 | 0.368 | 0.168 | 0.256 | -0.056 |
| 56 | Et | 3-OMe | 0.210±0.020 | 0.458 | 0.255 | -0.203 | 0.448 | 0.010 |
| 57 | iBu | 3-OMe | 0.400±0.110 | 0.632 | 0.213 | -0.419 | 0.605 | 0.027 |
| 58* | Me | 3-Me | 0.090±0.010 | 0.300 | 0.556 | 0.256 | 0.415 | -0.115 |
| 59 | Me | 3-EtO | 0.230±0.010 | 0.480 | 0.606 | 0.127 | 0.627 | -0.147 |
| 60 | Me | 3-acetamido | 0.059±0.002 | 0.243 | 0.500 | 0.257 | 0.383 | -0.140 |
| 61 | Me | 3,5-dimethoxy | 0.063±0.001 | 0.251 | 0.480 | 0.229 | 0.503 | -0.252 |
| 62 | Me | 3,4,5-trimethoxy | 1.300±0.220 | 1.140 | 0.453 | -0.687 | 0.802 | 0.338 |
| 63 | Me | 4-Cl-3-OMe | 0.063±0.003 | 0.251 | 0.906 | 0.655 | 0.262 | -0.011 |

| Number | R<sup>3</sup> | IC<sub>50</sub> | sqrtIC<sub>50</sub> | HM |  | GBR |  |
| --- | --- | --- | --- | --- | --- | --- | --- |
|  |  |  |  | predict | residual | predict | residual |
| 64 |  | 0.052±0.006 | 0.228 | 0.638 | 0.410 | 0.597 | -0.369 |
| 65 |  | 0.180±0.030 | 0.424 | 0.538 | 0.114 | 0.415 | 0.009 |
| 66 |  | 0.150±0.010 | 0.387 | 0.532 | 0.145 | 0.483 | -0.096 |
| 67* |  | 0.140±0.020 | 0.374 | 0.380 | 0.006 | 0.256 | 0.119 |
| 68* |  | 0.230±0.010 | 0.480 | 0.533 | 0.053 | 0.433 | 0.046 |
| 69 |  | 0.470±0.060 | 0.686 | 0.683 | -0.003 | 0.691 | -0.005 |
| 70 |  | 0.140±0.020 | 0.374 | 0.513 | 0.139 | 0.383 | -0.008 |
| 71 |  | 0.071±0.004 | 0.266 | 0.663 | 0.397 | 0.597 | -0.331 |
| 72 |  | 0.770±0.030 | 0.877 | 0.475 | -0.403 | 0.503 | 0.374 |
| 73* |  | 0.085±0.011 | 0.292 | 0.473 | 0.182 | 0.503 | -0.212 |
| 74 |  | 0.047±0.020 | 0.217 | 0.681 | 0.464 | 0.691 | -0.474 |
| 75 |  | 0.056±0.002 | 0.237 | 0.531 | 0.294 | 0.433 | -0.197 |
| 76 |  | 0.940±0.020 | 0.970 | 0.583 | -0.387 | 0.652 | 0.318 |
| 77 |  | 0.220±0.080 | 0.469 | 0.675 | 0.206 | 0.691 | -0.222 |
| 78 |  | 0.130±0.030 | 0.361 | 0.342 | -0.018 | 0.362 | -0.002 |
| 79 |  | 0.590±0.260 | 0.768 | 0.688 | -0.081 | 0.648 | 0.120 |
| 80 |  | 0.540±0.080 | 0.735 | 0.621 | -0.114 | 0.627 | 0.108 |
| 81 |  | 0.670±0.110 | 0.819 | 0.531 | -0.287 | 0.783 | 0.036 |

| Number | Structure | IC<sub>50</sub> | sqrtIC<sub>50</sub> | HM |  | GBRt |  |
| --- | --- | --- | --- | --- | --- | --- | --- |
|  |  |  |  | predict | residual | predict | residual |
| 82 |  | 0.100±0.010 | 0.316 | 0.363 | -0.047 | 0.362 | -0.046 |
| 83* |  | 0.027±0.003 | 0.164 | 0.178 | -0.014 | 0.195 | -0.031 |
| 84 |  | 0.710±0.020 | 0.843 | 0.677 | 0.166 | 0.802 | 0.040 |

sqrtIC<sub>50</sub>: The square root of IC<sub>50</sub>

### 2.2 Generation of descriptors

First of all, all of the 84 compound’s 2D structure were simply sketched by the software ChemDraw Ultra 8.0[11], and were saved as the mol file. Then, these compounds were inputted into HyperChem professional[12] in order to get pre-optimization by ways of MM+ and semi-empirical methods. After all steps were finished, we could get 4 file formats involving .mol, .mno, .hin and .zmt. Moreover, the MOPAC [13] was employed for geometrical optimization. Last but not least, the .mon and .zmt files were imported into the application CODESSA so that the descriptors of derivatives could be generated. The characters of descriptors in CODESSA were in abundance, such as geometrical descriptors, structural descriptors, topological descriptors and quantum descriptors [14].

### 2.3 The HM linear regression model

After the generation of descriptors, we fitted the HM linear regression model in CODESSA according to the datasets of descriptors. We selected several descriptors through testing cross-validated R^2^ (R^2^_cv_), coefficient of determination (R^2^), the standard deviation of error (S^2^) and the F test.

The strengths of HM model were obvious. Firstly, as a linear model, it was not only easy to model, but also having high interpretability[15]. Secondly, it had excellent advantages, it didn’t have software restriction and could obtain vintage model[16].

However, the result of fitting outcome was unsatisfactory, which proved that the relationship between descriptors and IC_50_ was complicated, therefore, a nonlinear model was established.

### 2.4 The GBR nonlinear model

In order to obtain a better result, we outputted two selected descriptors by means of python, we fitted these data to various models in python, such as support vector machine[17], random forest[18], gradient boosting regression and so on. Compared with other machine learning models, GBR showed wonderful fitting effect which could not be beat.

The main idea of GBR could be summarized as follows: add new models sequentially to the integration, a new weak-base learner model would be generated according to error of entire ensemble learned so far iteratively. As a framework, boosting could improve any wear-learning model when it came to every specific iteration, the error rate of weak model was only slightly better by contrast with random guessing, the remaining errors would be slightly improved by building up a simple weak model sequential model every sequential model. The basic algorithm for GBR could be generalized as follows, from which we could easily summerize that the finial model was just a stage additive model of b.

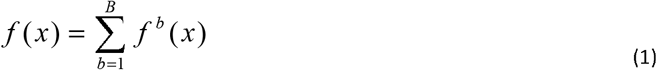

To get a better insight of GBR, a brief flowchart of GBR is given in Fig.1 and we will illustrate its algorithm flow, P is on behalf of parameter, which also includes multiply paramete. P = {p_0_, p_1_, p_2._…}, F (X; P) represents the function of X with P as the parameter, which is our predication function, several models combine to gain a better model, β represents the weight of each model, and α represents the parameters in the model. In order to optimize F, we can optimize {β, α} which is also named P.

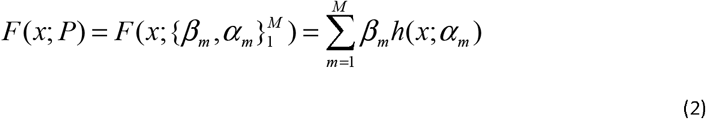

Φ(P) represents the likelihood function which is also the loss function of P:

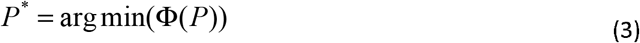

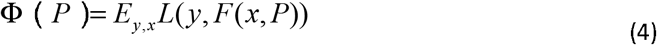

**Figure 1.**
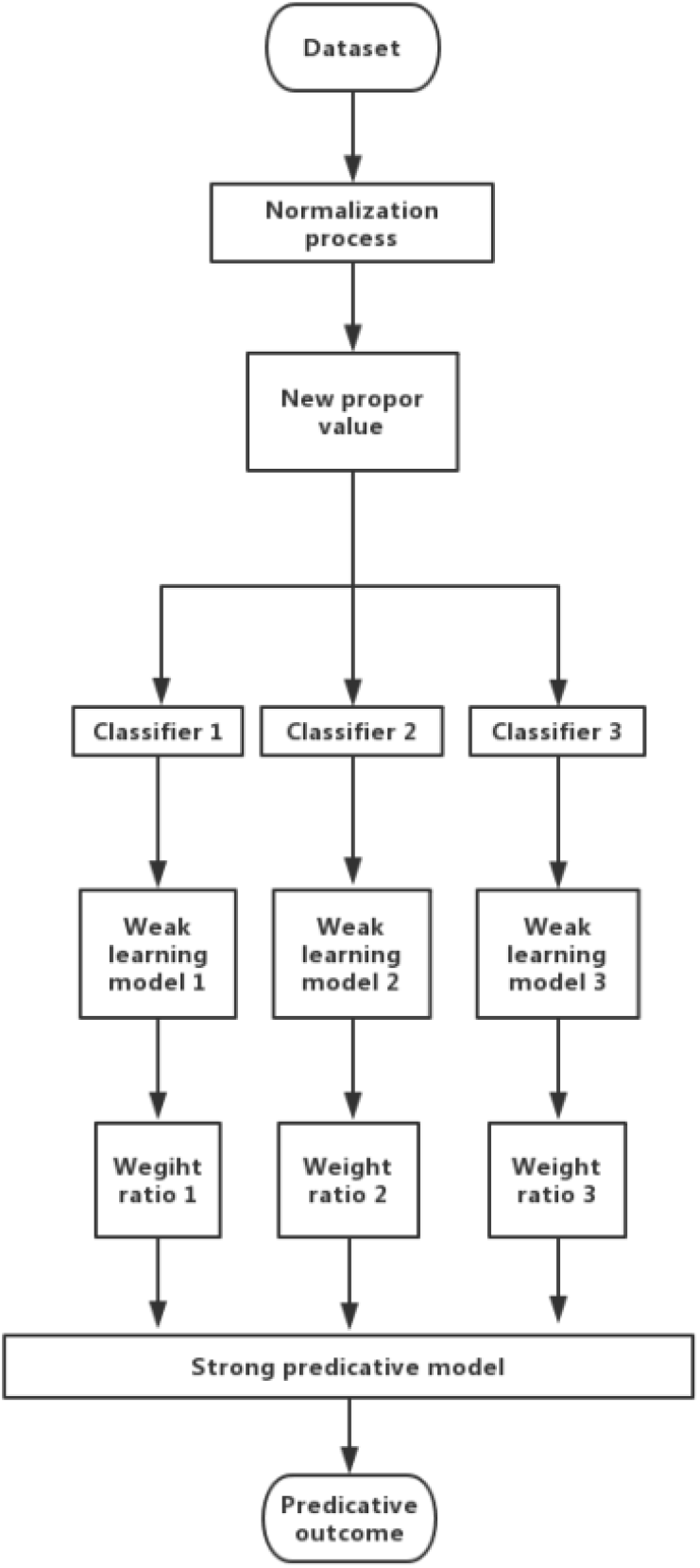
The Flowchart of GBR.

Since the model (F (x; P)) is additive, for the parameter P, we can also get the following formula:

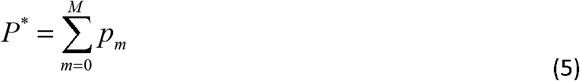

In this way, we consider the process of optimizing P as the process of gradient descent, assuming that m-1 models have been obtained, when we want to obtain the m-th model, we ought to get the gradient of the first m-1 models. g_m_ is the direction of the fastest decline.

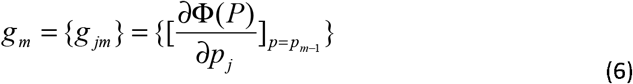

We assume that the first m-1 models are known, and we should never change these models, our concentration should be focus on the model established later:

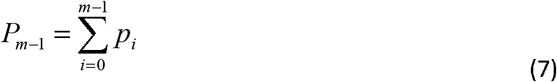

The new model we established is in the direction of the gradient of the P likelihood function, ρ is the descending distance in the gradient direction.

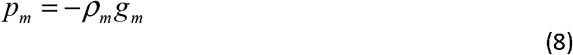

We can finally get the optimal ρ by optimizing the following formula:

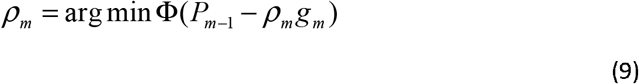

### 2.1 Comparison

In order to compare the results of these models, we calculated new statistical indicators of R^2^_CV_ and S^2^ with python. The indicators were calculated with the following equation:

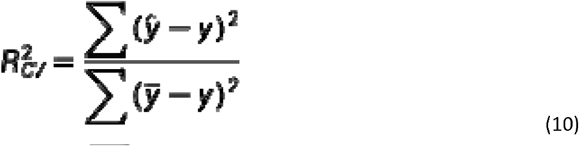

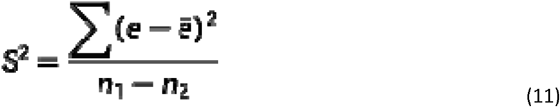

e: residual between observed value and predict value

n_1_: numbers of structures

n_2_: numbers of descriptors

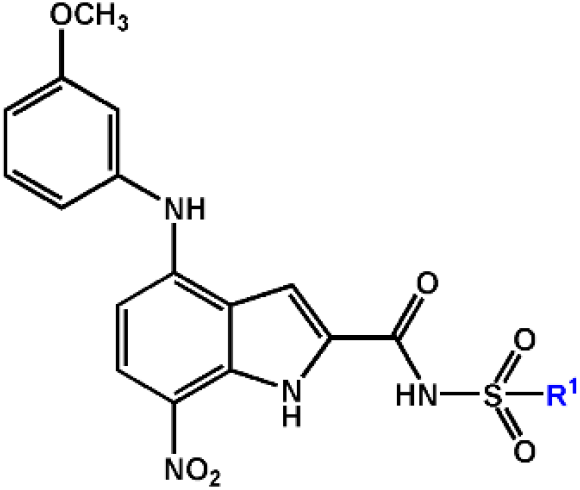

## 3. Results

### 3.1 Result of HM

618 descriptors for derivatives were analyzed by CODESSA program, the influence of the numbers of descriptor on the R^2^, R^2^_CV_ and S^2^ was showed in the Fig.2. We can see in the Fig.2 that as the numbers of descriptors improves, the R^2^, R^2^_CV_ grows, however, the S^2^ decreases. The change between one descriptor to two descriptors influences the values a lot, while after the number has changed to two, the growth of descriptors has little impact on data. Given that overfitting can be caused by excessive descriptors, two descriptors were selected to describe the activity of compounds. The correlation between two descriptors is shown in the Fig.3. In the Fig.3, the descriptor1 is on behalf of Min electroph react index for a C atom (MERICA) while the descriptor2 is on behalf of Min nucleoph react index for a S atom (MNRISA) and we can know that the correlation relationship between two descriptors is small, thus, there is no collinearity problem.

**Figure 2.**
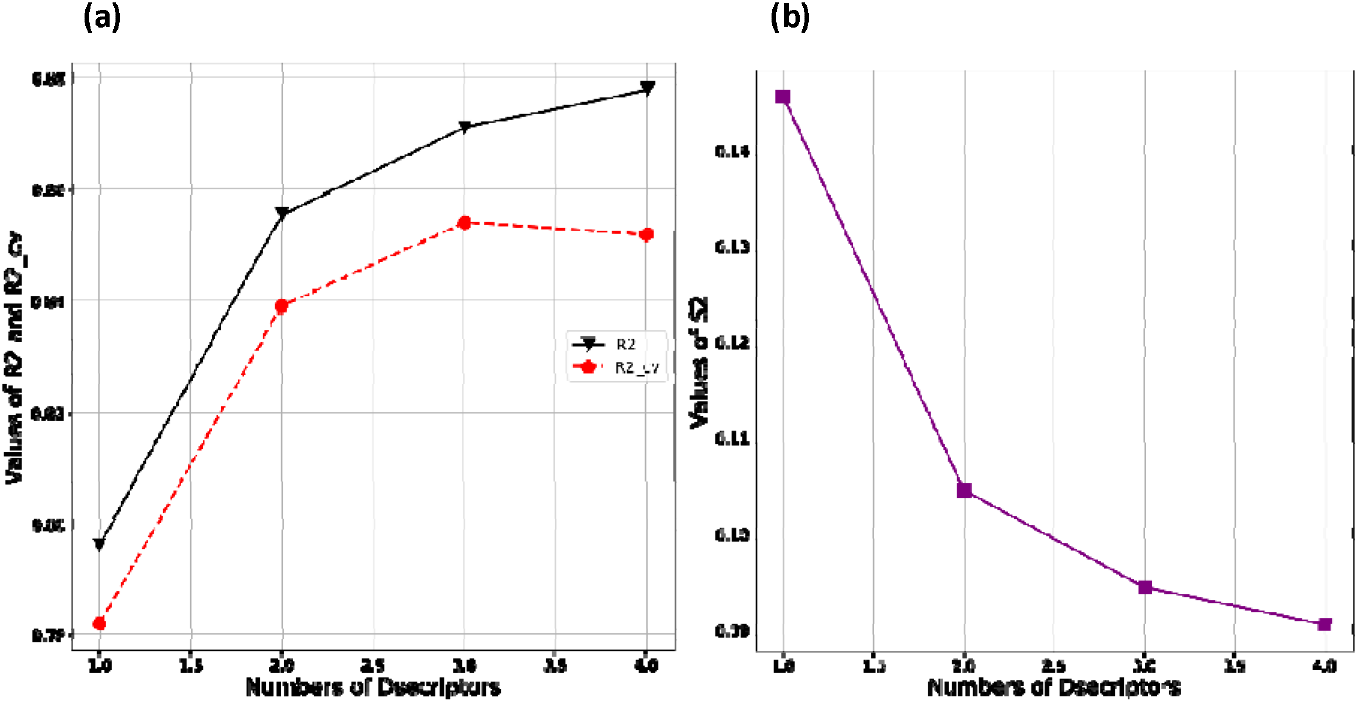
The Influence of the Numbers of Descriptor on R^2^, R^2^_CV_ and S^2^. (a) values of R^2^ and R^2^cv with increase of descriptors, (b)values of S^2^ with increase of descriptors

**Figure 3.**
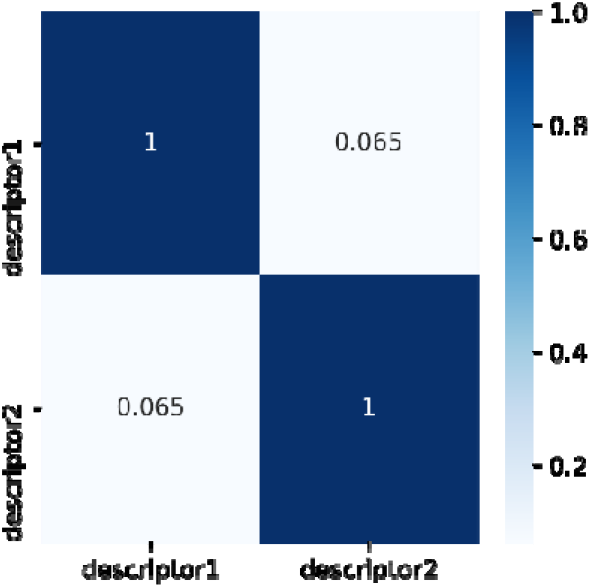
The Correlation Between Two Descriptors in the Model.

Tab.1 shows that the predicated outcome observed by HM based on 2 descriptors, the linear plot of HM is displayed in the Fig.4. The equation of QSAR model based on the HM is shown below.

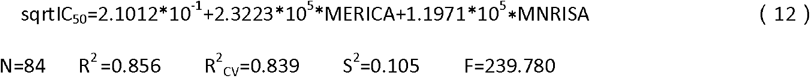

**Figure 4.**
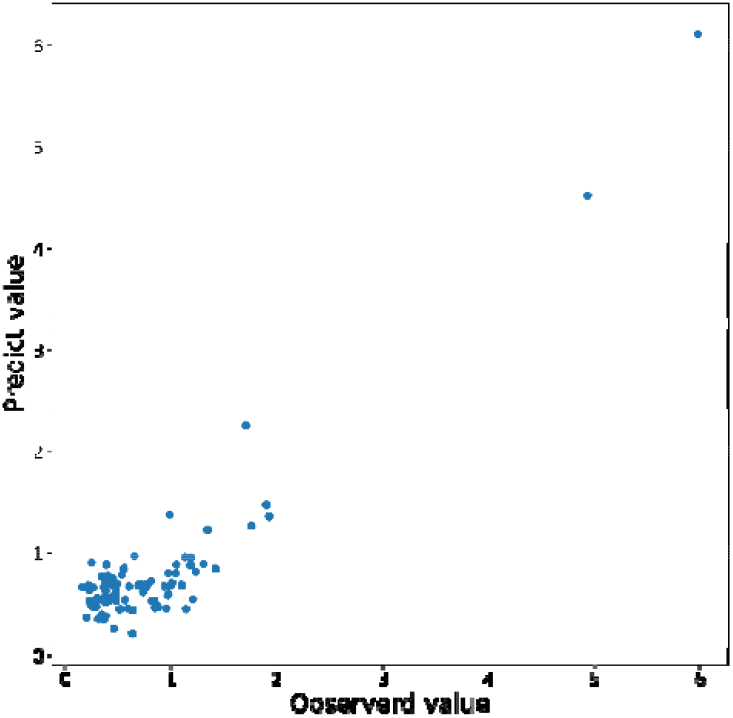
Linear Plot of Predicted Value versus Observed Value.

### 3.2 Result of GBR

Same descriptors and half-inhibitory concentration (IC_50_) were introduced into the nonlinear model GBR, which was written in python language. The predicated results are given in the Tab.1, GBR is a popular machine learning algorithm which has been proven to gain success in various field. We performed 2.7millions iterations, after which a good result with a R^2^ of 0.943 was achieved.

The detailed statistical results is displayed in the Tab.2. From Tab.2, we can see all R^2^ are bigger than 0.8, which means the model has a strong predictive ability. Fig.5 and Fig.6 shows the fitting curve of the training set and test set respectively and the frequency of residual is given in the Fig.7. It can be inferred from the Fig.5 and Fig.6 that the fitting effect of GBR model is excellent, and the error between the predicted value and the observed value is small. Compound 22, however, was somewhat less accurately predicted, with error of -0.776. It can be seen from the Fig.7 that the residuals predicted by GBR model approximately obey the standard normal distribution, which indicates that the residuals are independent of each other and there is no need to model the residuals. Meanwhile, most of the residual between predicted and observed value in GBR focus on the interval from -0.2 to 0.2, proving the predication accuracy of GBR.

**Table 2.**
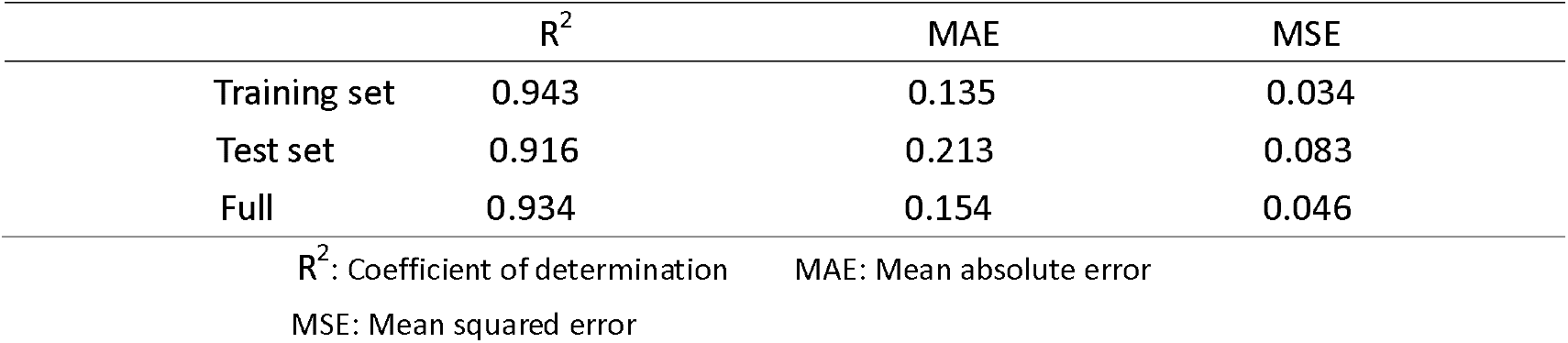
The Statistical Results of GBR Model.

**Figure 5.**
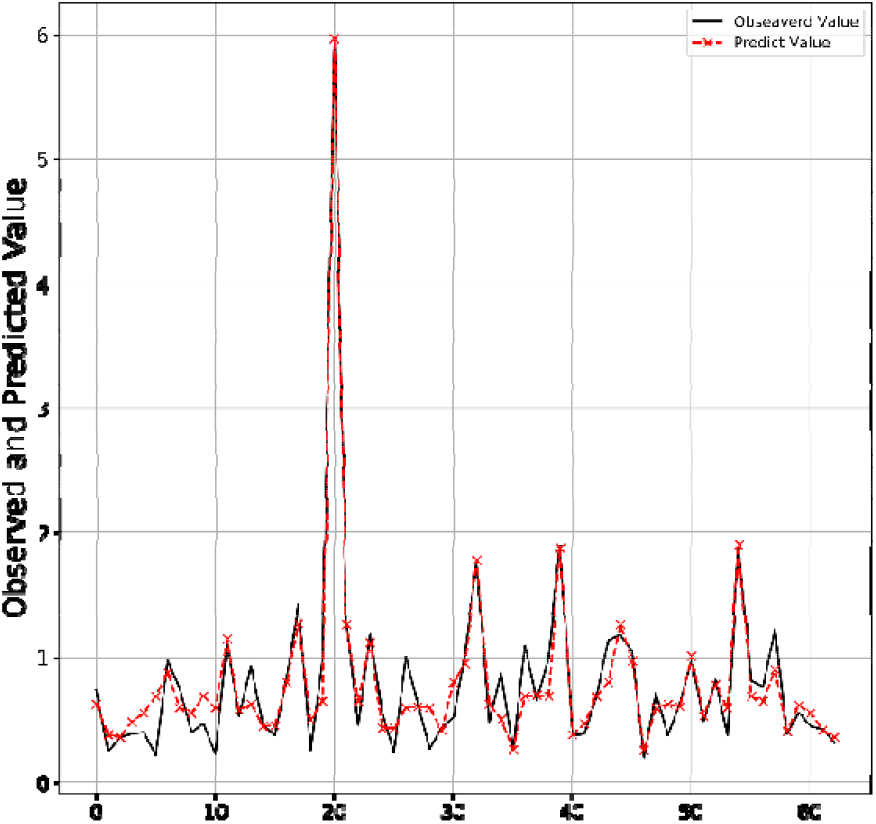
Fitting Curve of Training Set.

**Figure 6.**
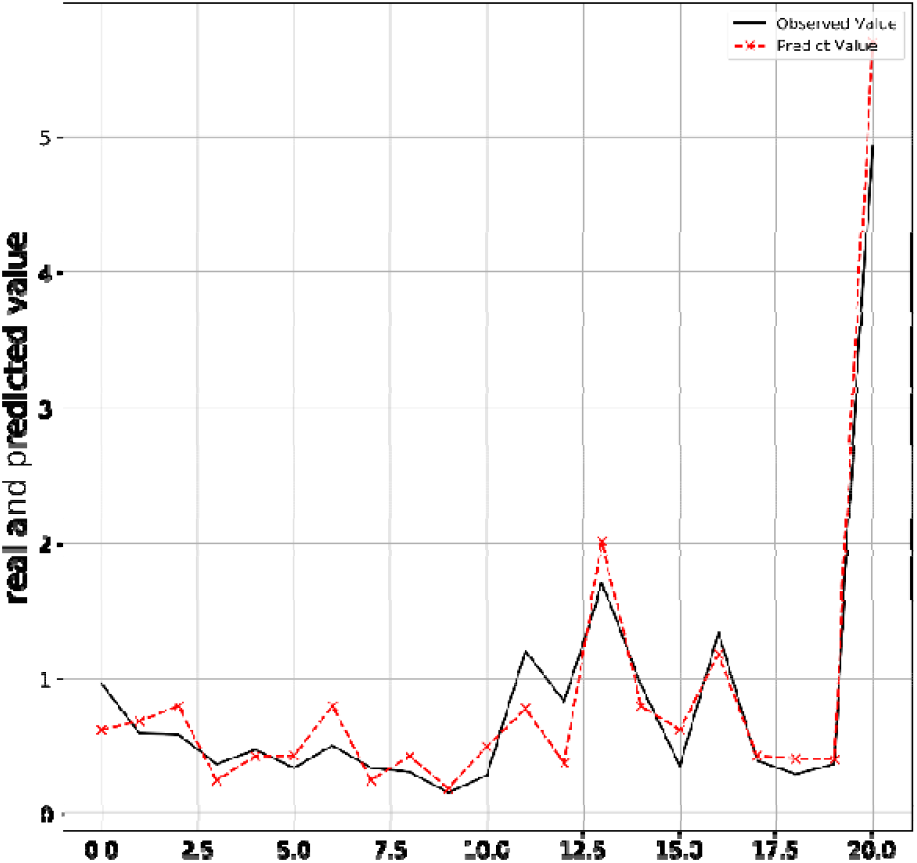
Fitting Curve of Test Set.

**Figure 7.**
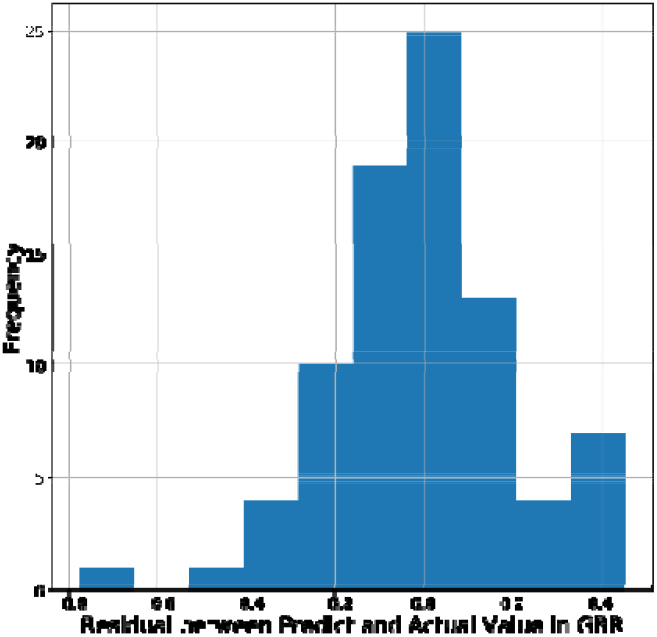
The Frequency Histogram of Residual in GBR.

### 3.3 Comparison

In order to compare the results of two models intuitively, we calculated new statistical evaluation indicators of two models, which is displayed in the Tab.3. Apparently, the R^2^, R^2^_CV_ of training set in GBR is much higher compared with the same index in HM, meanwhile, the S^2^ of the training set in GBR is smaller compared with HM. As a consequence, it is the nonlinear model GBR that demonstrates a better predicative effect.

**Table 3.** The common measure of HM and GBR.

| | $R^2$ | $R^2_{cv}$ | $S^2$ |
| --- | --- | --- | --- |
| HM | 0.856 | 0.839 | 0.105 |
| GBR | 0.943 | 0.925 | 0.046 |

$R^2$ : Coefficient of determination $R^2_{cv}$ : cross-validated coefficient of determination
$S^2$ : Standard Deviation of error

## 4. Discussion

### 4.1 The discussion of new GBR method applied in QSAR

GBR is frequently applied in the field concerning mathematical analysis and high energy physics, we noticed that few studies of drug design used this method of GBR. GBR could optimize different loss function and provides some hyperparameter adjustment option, making the function flexible, therefore, models generated by the GBR usually provide great predictive accuracy.

There is no denying that GBR has its own advantages. Firstly, the classification values and numerical values can be often applied well without any data pre-processing work. Secondly, there is no need to interpolate the loss data. Last but not least, it has excellent generalization ability because it utilizes the linear combination of multiple learners to gain the predicative accuracy, avoiding the problem of poor predicative effect caused by limited ability of single learner.

Coins have two sides, there is no doubts that the strengths of GBR is obvious. The shortages of GBR should not be ignored at the meantime. GBR pays much attention to outliers which leads to overfitting probably, so that cross-validation is of great importance to neutralize. The final model of the algorithm is obtained by integrating several sub-models, the high training speed will result in the ignorance of the sample information, which will also lead to overfit. Under this circumstance, we would better add more parameters to control the learning rate of the sub-algorithm model. Meanwhile, chances are that the high flexibility will affect the the behavior of the method (number of iterations, normalization parameters, ect). As a result, more studied ought to be done in order to minimize the bad influences of GBR towards the outcome.

### 4.2 The discussion of descriptors

It is important to avoid the collinearity in the process of developing of the multiple regression equation [19]. The correlation coefficient between descriptors is given in the Fig.3 with the good result of 0.065, which implies that the statistical relationship between descriptors is extremely small and proves the statistical reliability of the method.

In order to get a better understanding of the features that influence the activity of the derivative, we did further research on the chemical and physical function of the descriptors.

Min electroph.react.index for a C atom (MERICA) is a kind of electronic molecule descriptors, which has positive-sign coefficient according to the Eq.(12) in the HM. Obviously, this detail reflects that increasing MERICA enhance the IC50 of N-Arylsulfonyl-Indole-2-Carboxamide derivatives and indicates the quantum chemical description of C atom of of great significance. MNRISA is a quantum mechanical descriptors measuring Min nucleoph.react index for a S atom. Compared with MERICA, MNRISA also have positive-sign coefficient in the equation, however, its coefficient is much smaller than MERICA. Hence, it is convinced that the raise of MNRISA would directly increase the value of IC_50_, despite that, MERICA’s impact on IC_50_ is far higher than MNRISA. In addition, all compounds in this group involve the C and S atom in their structure, which emphasizthees the necessity of introducing these descriptors into the model. Essentially, the growth of MERICA and MNRISA implies that the nuclear and electronical reaction are more difficult to occur, thus, unstable ions and nuclei are less likely to be generated, which may attact FBPase to inhibit its activity. As a result, the increase of the descriptors suggests the inhibitory ability of coupounds will attenuate. In a word, the IC50 of the coupounds will increase.

## 5. Conclusion

We built linear model with HM in CODESSA software and nonlinear model with GBR method in python. We got the models of the inhibitory relationship between Fructose-1,6-Bisphosphatase and N-Arylsulfonyl-Indole-2-Carboxamide. This study makes a bold attempt at the application of new method GBR in QSAR and proves GBR is a promising tool for further study of CADD. In addition, our model displays how Min electroph.react.index for a C atom and Min nucleoph.react index for a S atom affect the bioactivity. Thus, the results of our study provide a useful guideline and support for potential new drugs of T2DM.

## Reference

1. Murphy, H.R., et al., Characteristics and outcomes of pregnant women with type 1 or type 2 diabetes: a 5-year national population-based cohort study. The Lancet Diabetes & Endocrinology, 2021. 9(3): p. 153–164.

2. Lim, S., et al., Diabetes drugs and stroke risk: Intensive versus conventional glucose-lowering strategies, and implications of recent cardiovascular outcome trials. Diabetes, Obesity and Metabolism, 2020. 22(1): p. 6–15.

3. Kaur, R., L. Dahiya, and M. Kumar, Fructose-1,6-bisphosphatase inhibitors: A new valid approach for management of type 2 diabetes mellitus. European Journal of Medicinal Chemistry, 2017. 141: p. 473–505.

4. Padhi, S., A.K. Nayak, and A. Behera, Type II diabetes mellitus: a review on recent drug based therapeutics. Biomedicine & Pharmacotherapy, 2020. 131: p. 110708.

5. Exton, J.H., Gluconeogenesis. Metabolism, 1972. 21(10): p. 945–990.

6. Chen, L., et al., Cloning, purification and characterisation of cytosolic fructose-1,6-bisphosphatase from mung bean (Vigna radiata). Food Chemistry, 2021. 347: p. 128973.

7. Barciszewski, J., et al., T-to-R switch of muscle fructose-1,6-bisphosphatase involves fundamental changes of secondary and quaternary structure. Acta Crystallographica Section D, 2016. 72(4): p. 536–550.

8. Zhao, L., et al., Advancing computer-aided drug discovery (CADD) by big data and data-driven machine learning modeling. Drug Discovery Today, 2020. 25(9): p. 1624–1638.

9. Cruz-Monteagudo, M., et al., Systemic QSAR and phenotypic virtual screening: chasing butterflies in drug discovery. Drug Discovery Today, 2017. 22(7): p. 994–1007.

10. Zhou, J., et al., Discovery of N-Arylsulfonyl-Indole-2-Carboxamide Derivatives as Potent, Selective, and Orally Bioavailable Fructose-1,6-Bisphosphatase Inhibitors—Design, Synthesis, In Vivo Glucose Lowering Effects, and X-ray Crystal Complex Analysis. Journal of Medicinal Chemistry, 2020. 63(18): p. 10307–10329.

11. Mendelsohn, L.D., ChemDraw 8 Ultra, Windows and Macintosh Versions. Journal of Chemical Information and Computer Sciences, 2004. 44(6): p. 2225–2226.

12. Froimowitz, M., HyperChem : a software package for computational chemistry and molecular modeling. BioTechniques, 1993. 14(6): p. 1010–1013.

13. Stewart, J.J.P., MOPAC: A semiempirical molecular orbital program. Journal of Computer-Aided Molecular Design, 1990. 4(1): p. 1–103.

14. Wang, Y., et al., Quantitative structure–activity relationship for prediction of the toxicity of polybrominated diphenyl ether (PBDE) congeners. Chemosphere, 2006. 64(4): p. 515–524.

15. Graybill, F. Theory and Application of the Linear Model. 1976.

16. Song, R., et al. QSAR study on the IC_(50) of 6-alkenylamides of 4-anilinothieno[2,3-d] pyrimidine as epidermal growth factor receptor inhibitors in lung cancer. 2015.

17. Noble, W.S., What is a support vector machine? Nature Biotechnology, 2006. 24(12): p. 1565–1567.

18. Pal, M., Random forest classifier for remote sensing classification. International Journal of Remote Sensing, 2005. 26(1): p. 217–222.

19. Si, H., et al., Predicting the activity of drugs for a group of imidazopyridine anticoccidial compounds. European Journal of Medicinal Chemistry, 2009. 44(10): p. 4044–4050.

